# Genomic resources for *Goniozus legneri*, *Aleochara bilineata* and *Paykullia maculata*, representing three independent origins of the parasitoid lifestyle in insects

**DOI:** 10.1101/300418

**Authors:** Ken Kraaijeveld, Peter Neleman, Janine Mariën, Emile de Meijer, Jacintha Ellers

## Abstract

Parasitoid insects are important model systems for a multitude of biological research topics and widely used as biological control agents against insect pests. While the parasitoid lifestyle has evolved numerous times in different insect groups, research has focused almost exclusively on Hymenoptera from the parasitica clade. The genomes of several members of this group have been sequenced, but no genomic resources are available from any of the other, independent evolutionary origins of the parasitoid lifestyle. Our aim here was to develop genomic resources for three parasitoid insects outside the parasitica. We present draft genome assemblies for *Goniozus legneri*, a parasitoid Hymenopteran more closely related to the non-parasitoid wasps and bees than to the parasitica wasps, the Coleopteran parasitoid *Aleochara bilineata* and the Dipteran parasitoid *Paykullia maculata.* The genome assemblies are fragmented, but complete in terms of gene content. We also provide preliminary structural annotations. We anticipate that these genomic resources will be valuable for testing the generality of findings obtained from parasitica wasps in future comparative studies.

**Data availability:** The Whole Genome Shotgun projects have been deposited at DDBJ/ENA/GenBank under the accessions NCVS00000000 (*G. legneri*), NBZA00000000 (*A. bilineata*) and NDXZ00000000 (*P. maculata).* The versions described in this paper are versions NCVS01000000, NBZA01000000 and NDXZ01000000, respectively. Mapped reads and genome annotations are available through http://parasitoids.labs.vu.nl/parasitoids/. This website also includes genome browsers and viroblast instances for each genome.

## Introduction

Parasitoid insects have long been used as model systems for the study of a wide variety of topics in biology, including life history, chemical ecology and population dynamics (Godfray 1994; Hawkins and Sheehan 1994; Wajnberg and Colazza 2013). Parasitoids are also widely employed as agents of biological control against insect pests (Heimpel and Mills 2017). In recent years, the study of parasitoid insects has received new impetus through the availability of a steadily growing number of available genome sequences. Genomes of 13 parasitoid insects have recently become available, all from within one monophyletic clade of parasitoid wasps (Branstetter *et al.* 2018). These genomes are rapidly becoming a rich source of information on many aspects of parasitoid biology, e.g. (Werren *et al.* 2010).

The vast literature on insect parasitoids deals almost exclusively with Hymenopteran parasitoids, which all share a single evolutionary origin (Eggleton and Belshaw 2013). The stinging wasps (Aculeata) diverged from this group and lost the parasitoid life style. However, recent phylogenetic studies place the parasitoid Chrysidoidea within the Aculeata, suggesting that they may have re-evolved the parasitoid life style secondarily (Carr *et al.* 2010; Peters *et al.* 2017). Outside the Hymenoptera, parasitoid lifestyles have evolved in multiple insect groups, including Diptera, Coleoptera, Lepidoptera, and Neuroptera (Eggleton and Belshaw 2013). Dozens, or even hundreds, of evolutionarily independent parasitoid lineages are found within the Diptera and Coleoptera (Feener and Brown 1997; Eggleton and Belshaw 2013). It has been estimated that 20% of all parasitoid insect species are Dipterans (Feener and Brown 1997). Some of these are economically important, either as biological control agent (Grenier 1988) or as pest (Prabhakar *et al.* 2017). The study of such non-hymenopteran parasitoid systems would benefit from genomic resources, as it is unknown to what extent insights from hymenopteran parasitoids can be extrapolated to non-hymenopteran parasitoids. Unfortunately, no sequenced genomes are available for any of these groups as yet.

Here, we present draft genome assemblies for three parasitoid insect species that each represent an evolutionary independent acquisition of the parasitoid lifestyle (Figure 1A). *Goniozus legneri* is a parasitoid wasp from the superfamily Chrysidoidea (family Bethylidae) that is not part of the species-rich and well-studied parasitica clade. *G. legneri* can therefore function as an outgroup in comparative analyses of parasitica wasps and may have re-evolved the parasitoid lifestyle after it was lost at the base of the Aculeata. *G. legneri* is a gregarious parasitoid of Lepidopteran larvae that stings and paralyzes its prey before ovipositing on it externally (Khidr *et al.* 2013). The female then guards the host against utilization by other females (Khidr *et al.* 2013). *Aleochara bilineata* is a Coleopteran parasitoid of Dipteran pupae that represents another evolutionary independent acquisition of the parasitoid lifestyle. Females lay their eggs in the proximity of hosts (Bili *et al.* 2016). After hatching, the first instar larvae search for, enter and parasitize host pupae (Bili *et al.* 2016). *Paykullia maculata* is a parasitoid fly from the family of the Rhinophoridae, representing one of the many independent acquisitions of the parasitoid mode within the Diptera. Like all Rhinophoridae, *P. maculata* parasitizes isopods. Females lay their eggs in the vicinity of isopod aggregations. The larvae latch on to a passing isopod and enter it through an intersegmental membrane (Wijnhoven 2001). They feed on the isopod’s hemolymph and later also on non-vital tissues, like the female ovaria (Wijnhoven 2001). The isopod continues to feed and moult normally, until the parasitoid kills it and pupates within the host exoskeleton (Wijnhoven 2001). Here, we present draft genome sequences for these three species.

**Figure 1.**
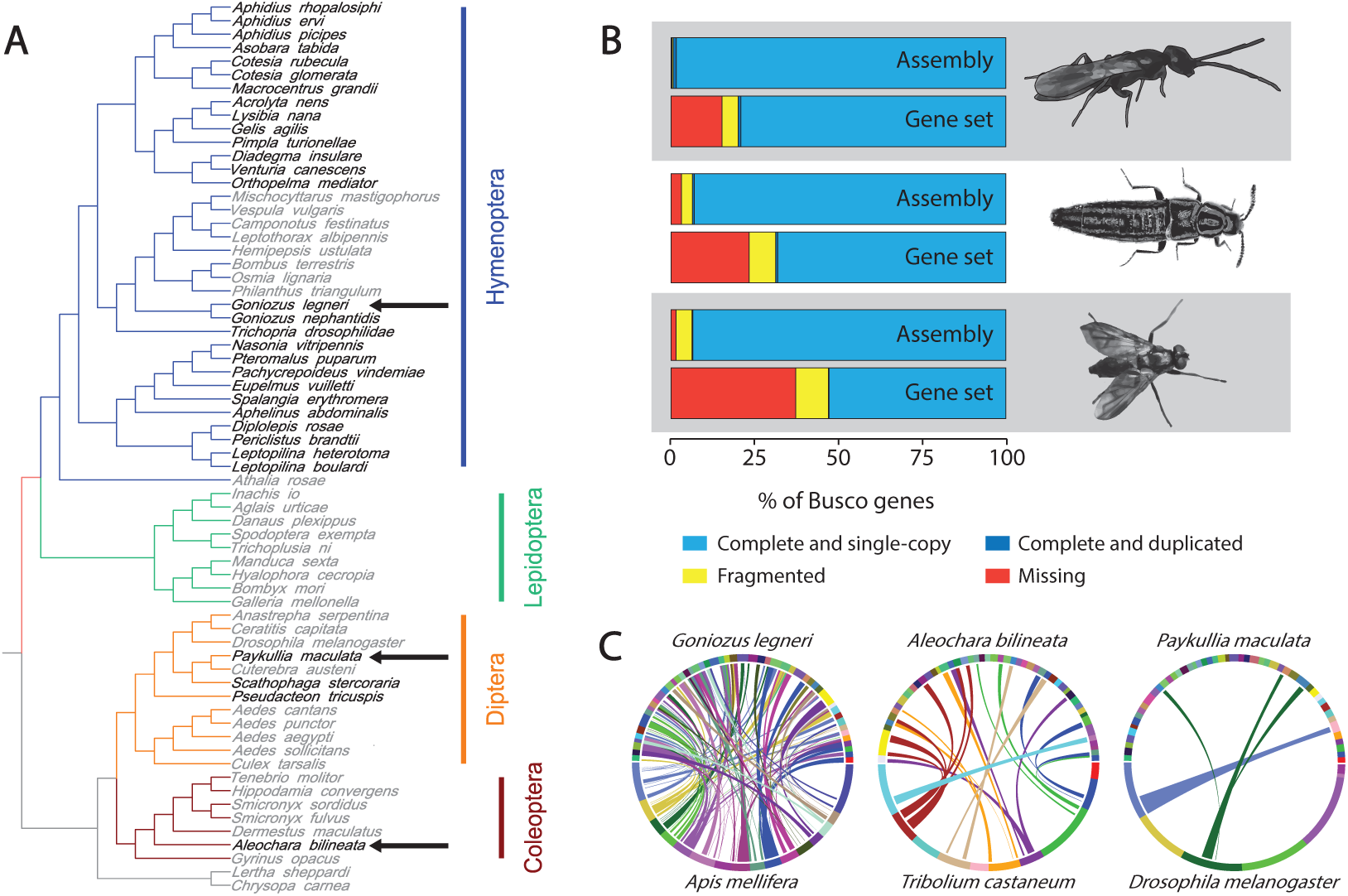
Phylogenetic context and genome features. (A) Phylogeny of selected insect species from (Visser *et al.* 2010). Parasitoid species are indicated in black font and non-parasitoid species in grey font. The three species for which we provide draft genome assemblies are indicated by arrows. (B) Completeness of the genome assemblies and annotation. Orthologs of 1658 genes that are present as single copies in at least 90% of insects were retrieved using Busco v3. From top to bottom: *Goniozus legneri* genome, *G. legneri* gene set, *Aleochara bilineata* genome, *A. bilineata* gene set, *Paykullia maculata* genome and *P. maculata* gene set. (C) Circle plots illustrating the synteny between the 50 largest scaffolds of the genome assemblies of *G. legneri, A. bilineata* and *P. maculata* (top half of each circle) and the genomes of the closest available relative with chromosome- or linkage group-level assemblies (A. *mellifera, T. castaneum* and *D. melanogaster*, respectively; bottom half of each circle). Links are colored to match the chromosome or linkage group of the model species.

## Materials and Methods

### Samples

Specimens of *G. legneri* were obtained from a long-term culture at the University of Nottingham (UK), which is descended from founders originally collected in Uruguay (I. Hardy, pers. comm.). Specimens of *A. bilineata* were obtained from a culture maintained at the University of Rennes (France). Specimens of *P. maculata* were cultured from *Porcellio scaber*, collected in the field in The Netherlands (Wageningen and Amsterdam) during 2015. For each species, DNA was extracted from a single adult female using different spin column protocols. For *G. legneri* and *A. bilineata*, DNA was obtained by crushing the insect in 100-200 μl PBS and adding 100 μl nuclei lysis buffer, 4 μl Proteinase K, 5 μl RNAse (all Promega) and incubating at 60°C for 15 min. A further 340 μl lysis buffer was then added and the sample was centrifuged at full speed for 5 min. The supernatant was then transferred to a spin column (Promega), rinsed with 500 μl wash buffer (Promega) three times and eluted in 100 μl H20. *P. maculata* was crushed in liquid nitrogen and DNA was extracted using QIAamp DNA Mini Kit (Qiagen) following the manufacturer’s protocol. The quantity and quality of the DNA samples was assessed using Nanodrop en Qubit.

### Whole genome sequencing

Two Truseq fragment libraries with slightly different insert sizes were constructed from each DNA sample. The two libraries for each species were barcoded, pooled and sequenced using 2×100 bp paired-end sequencing on Illumina HiSeq 2000 (*G. legneri* and *A. bilineata*), or 2×125 bp paired-end on Illumina HiSeq 2500 (*P. maculata*).

### De novo genome assembly

Prior to *de novo* assembly, we characterized the raw read data using SGA (Simpson 2014) and estimated genome size using KmerGenie (Chikhi and Medvedev 2014). Furthermore, we removed co-sequenced genomes to reduce complexity of the read set. To this end, reads were mapped to the mitochondrial DNA of well-sequenced related species (*Apis mellifera*, *Tribolium castaneum* and *Drosophila melanogaster* for *G. legneri*, *A. bilineata* and *P. maculata*, respectively) and to a panel of 12 *Wolbachia* strains (*w*Au GCA_000953315.1; *w*Bm GCA_000008385.1; *w*Cle GCA_000829315.1; *w*Pip_Pel GCA_000073005.1; *w*Ha GCA_000376605.1; *w*No GCA_000376585.1; *w*Ri GCA_000022285.1; *w*Mel GCA_000008025.1; *w*Fol CP015510.1; *w*Oo GCA_000306885.1; *w*Ov GFA_000530755.1; *w*Pip GCA_000208785.1) using Bowtie2 (Langmead and Salzberg 2012) with default settings. Unmapped reads were extracted from the Sam file using samtools view with -f 4 (Li *et al.* 2009) and reverted back to fastq using bedtools bamToFastq (Quinlan and Hall 2010). Reads were trimmed using platanus_trim (Kajitani *et al.* 2014). Given that the results from the SGA analysis indicated high levels of heterozygosity for *A. bilineata* and *P. maculata*, we chose to perform the *de novo* assembly in Platanus, which is specifically geared to deal with short-read data from heterozygous genomes (Kajitani *et al.* 2014). Assembly was followed by scaffold and gap_close steps as implemented in the Platanus pipeline. To assess coverage, reads were mapped back to the assembly using Bowtie2 with default settings and coverage was calculated using bedtools.

### Annotation

Structural annotation was performed using Maker2 using default settings (Holt and Yandell 2011). Annotation statistics were obtained using GAG (Hall *et al.* 2014). The completeness of the genomes and gene sets was assessed by identifying the number of insect Benchmark Universal Single-Copy Ortholgs (BUSCOs) (Simao *et al.* 2015). BUSCO v3.0.2 was run on both the genome assembly (using -m geno) and the Maker gene set at the peptide level (using -m prot) with the insecta_odb9 lineage dataset as reference. We compared the draft genome of each species to that of its closest available relative with a chromosome- or linkage group-level genome sequence available (*Apis mellifera* Amel_4.5, *Tribolium castaneum* Tcas5.2 and *Drosophila melanogaster* release 6 for *G. legneri*, *A. bilineata* and *P. maculata*, respectively) using SyMap v4.2 (Soderlund *et al.* 2011).

### Co-sequenced genomes

To remove any co-sequenced genomes (in addition to *Wolbachia* and mitochondria, which were removed prior to assembly), we employed the Blobology pipeline (Kumar *et al.* 2013).

## Results and Discussion

We generated 18.6-43.6 Gb data per species, covering each genome >100× (Table 1). These data were assembled into draft genomes that were reasonably close to the predicted size for each species (Table 2). The genome size of *G. legneri* appears small compared to other sequenced genomes of Hymenoptera, but is within the range for parasitoid wasps (e.g. *Macrocentrus cingulum*: 128 Mb, *Fopius arisanus*: 153 Mb). The estimated genome size of *A. bilineata* is smaller than that of other sequenced Coleoptera (smallest to date is *Hypothenemus hampei:* 151 Mb). The genome size of *P. maculata* is within the range observed for Diptera (e.g. *Zaprionus indianus* 124 Mb; *Rhagoletis zephyria:* 795 Mb). Further work is required to establish the accuracy of our genome size estimates.

**Table 1.** Raw reads generated for assembly.

| Species | Read pairs | Base pairs aligned | Coverage |
| --- | --- | --- | --- |
| *Goniozus legneri* | 140.0 M | 22.52 Gb | 160.8x |
| *Aleochara bilineata* | 170.3 M | 18.55 Gb | 217.8x |
| *Paykullia maculata* | 211.9 M | 43.56 Gb | 103.4x |

**Table 2.** Assembly summary statistics.

| Metric | *G. legneri* | *A. bilineata* | *P. maculata* |
| --- | --- | --- | --- |
| GC (%) | 40.6 | 39.9 | 28.4 |
| Scaffold count | 7,863 | 33,003 | 147,656 |
| Contig count | 13,705 | 40,228 | 169,825 |
| Total length (Mb) | 140.1 | 85.9 | 422.4 |
| Predicted length KmerGenie (Mb) | 142 | 112 | 536 |
| Gap (%) | 0.18 | 1.1 | 0.29 |
| Scaffold N50 (kb) | 167.3 | 54.1 | 7.7 |
| Contig N50 (kb) | 37.8 | 12.1 | 5.9 |
| Max. scaffold length (Mb) | 1.5 | 0,876 | 0.17 |
| Max. contig length (kb) | 371.7 | 316.8 | 86.6 |
| Number of scaffolds > 50 kb | 684 | 445 | 108 |
| % of genome in scaffolds > 50 kb | 82.4 | 52.1 | 1.8 |

The genome assemblies presented here were obtained from outbred, heterozygous individuals. The Platanus assembler is specifically designed to handle such data and creates a mosaic of the two haplotypes (Kajitani *et al.* 2014). We tested this by mapping the sequence reads back to the genome assembly. Bowtie2 in default settings reports only the best alignment for each read and haplotypes that had assembled as separate contigs should have half the coverage as collapsed haplotypes, as reads would only map to one of the haplotypes. In our case, the coverage histograms were unimodal, indicating that haplotypes were successfully collapsed.

The assembly for *G. legneri* yielded hits to Rhabditida nematodes, some of which are known parasites of insects. A Blast search of the entomopathogenic Rhabditidid *Oscheius sp.* (Lephoto *et al.* 2016) against the *G. legneri* genome assembly revealed two hits, upon which we removed one contig and one partial contig from the assembly. *A. bilineata* contained *Wolbachia*, but no other co-sequenced genomes. *P. maculata* contained no co-sequenced genomes.

The genome assemblies are very complete, with 97-99% of BUSCOs present and only 0.4-4.7% fragmented (Figure 1B). Lacking transcriptome data and other genomic resources for these or closely related species, the structural annotation is less complete. Maker2 annotated 5588-7463 genes per genome (Table 3), which is below the expected value for eukaryotes. BUSCO analysis indicated the gene sets to be 63-85% complete (Figure 1bB). It is thus imperative for future studies to interrogate the genome assembly for genes missing from the gene set.

**Table 3.** Gene annotation summary statistics.

| Metric | *A. bilineata* | *G. legneri* | *P. maculata* |
| --- | --- | --- | --- |
| Number of genes | 7220 | 7463 | 5588 |
| Number of exons | 28720 | 46322 | 20856 |
| Total gene length (Mb) | 17.2 | 25.5 | 18.1 |
| Mean gene length | 2380 | 3418 | 3232 |
| % of genome covered by<br>genes | 20.0 | 18.2 | 4.3 |
| mean exons per mRNA | 4 | 6 | 4 |

The level of synteny to well-characterized genome sequences of related model species varied (Figure 1C). The draft genome of *G. legneri* shows many collinear regions with the genome of *A. mellifera*, while the similarity between *P. maculata* and *D. melanogaster* is limited, with intermediate collinearity between *A. bilineata* and *T. castaneum* (Figure 1C). These differences are probably caused by a combination of factors, including quality of the draft genome assemblies, levels of relatedness to the selected model organism and rates of genome evolution.

In summary, we present fragmented, but relatively complete genome assemblies of three parasitoid insects, representing three independent evolutionary origins of the parasitoid lifestyle. These genomes will be valuable for comparisons to the widely studied parasitoid Hymenoptera, for which numerous genomes are available (Branstetter *et al.* 2018). Our study highlights that useful genomic resources can now be obtained from highly heterozygous individual insects collected from outbred lab cultures or even from the field, relieving the need for labor-intensive inbreeding procedures.

## Acknowledgements

We thank Ian Hardy, Denis Poinsot, Mikael Bili, Mark Lammers and Rik Zoomer for providing insect samples.

